# Thermally moderated firefly activity is delayed by precipitation extremes

**DOI:** 10.1101/074633

**Authors:** Sara L. Hermann, Saisi Xue, Logan Rowe, Elizabeth Davidson-Lowe, Andrew Myers, Bahodir Eshchanov, Christie A. Bahlai

**Affiliations:** Department of Entomology, Michigan State University; Department of Entomology, Pennsylvania State University; Program in Ecology, Evolutionary Biology and Behavior, Michigan State University; Biomass Conversion Research Laboratory, Department of Chemical Engineering, Michigan State University; DOE Great Lakes Bioenergy Research Center, East Lansing, Ml; Department of Integrative Biology, Michigan State University; Mozilla Science Lab

**Keywords:** Lightning bug, lampyridae, phenology, ecoinformatics, LTER

## Abstract

The timing of events in the life history of temperate insects is most typically primarily cued by one of two drivers: photoperiod or temperature accumulation over the growing season. However, an insect’s phenology can also be moderated by other drivers like rainfall or the phenology of its host plants. When multiple drivers of phenology interact, there is greater potential for phenological asynchronies to arise between an organism and those with which it interacts. We examined the phenological patterns of a highly seasonal group of fireflies (*Photinus spp*, predominantly *P. pyralis*) over a 12-year period (2004–2015) across 10 plant communities to determine if interacting drivers could explain the variability observed in the adult flight activity density (*i.e.* mating season) of this species. We found that temperature accumulation was the primary driver of phenology, with activity peaks usually occurring at a temperature accumulation of ~800 degree days (base 10°C), however, our model found this peak varied by nearly 180 degree day units among years. This variation could be explained by a quadratic relationship with the accumulation of precipitation in the growing season; in years with either high and low precipitation extremes at our study site, flight activity was delayed. More fireflies were captured in general in herbaceous plant communities with minimal soil disturbance (alfalfa and no-till field crop rotations), but only weak interactions occurred between within-season responses to climatic variables and plant community. The interaction we observed between temperature and precipitation accumulation suggests that, although climate warming has potential to disrupt phenology of many organisms, changes to regional precipitation patterns can magnify these disruptions.

## Introduction

Much can be learned about biological systems by observation alone [1], and observational data are often captured incidentally as a result of human activity [2], Incidental data can range from the very informal and uncontrolled (*e.g.* comments on a topic in a Web forum) to highly controlled and meticulously collected (*e.g.* unused data from scientific experiments). Indeed, research activities can produce systemic observational data of very high quality; for instance, insect trapping systems seldom only capture target taxa. This ’by-catch’ can provide data that support investigations into entirely uninvestigated phenomena. In this study we examine one such ’by-catch’ data set: a 12-year time series of firefly observations in southwestern Michigan, for their responses to environmental and habitat conditions.

Over 2,000 species of firefly (Coleoptera: Lampyridae) have been identified across various temperate and tropical environments around the world [3]. As larvae, species within the family Lampyridae spend much of their time living underground feeding on earthworms, mollusks, and other subterranean invertebrates [4], As adults, most species abstain from feeding [5], with the exception of the species *Photuris pennsylvanica*, of which the female is a voracious predator of both con-specifics as well as other insects [5–7], Few studies been conducted on firefly conservation and broader-scale ecology in relation to changing environments and land uses, and little is known about how environmental parameters drive firefly life history. It has been demonstrated that the life history of at least one species of firefly is temperature-dependent; researchers found that *P. pennsylvanica* adult emergence could be artificially accelerated by exposing larvae to increased soil temperature [8]. However, much of the primary research on fireflies has focused on the bio-luminescent properties of the firefly [9–14], while research describing basic population and community ecology of this important family is lacking.

In addition to the scientific importance of the Lampyridae for their bio-luminescent properties and as model organisms for evolutionary investigations, fireflies are also among the most widely recognized and culturally valued insect families among non-scientists. Two US states have designated a firefly as their “State Insect”[15]. Notably, fireflies also feature prominently in Japanese culture, where they have been designated as national natural treasures in many districts and have been used to generate support for biodiversity conservation efforts in Japanese agricultural regions [16–18]. They have also been touted as useful classroom tools for sparking student interest in biology [19]. Because of their popular appeal, it is unsurprising that public concern has grown about apparent declines of firefly populations from regions around the world where they occur [20].

Considering the paucity of ecological information about fireflies, their widespread popularity, and ease with which adults are observed, and concerns about their population viability, fireflies represent an ideal species for citizen science investigations. Citizen science efforts are currently underway seeking to gain information about the status, geographic distribution, and phenology of fireflies [21–23] and peer-reviewed publications on fireflies have already been produced based on these volunteer-generated data [24,25]. The popularity of fireflies gives them great potential as a flagship and umbrella conservation species and potentially an indicator species of ecological degradation in agricultural regions [26]. However, to our knowledge, no long-term systematic study of firefly phenology and responses to environmental drivers has been published.

Phenology plays a significant role in regulating species abundance, distribution, and biodiversity [27,28]. The timing of phenological events in insect life histories is strongly linked to climatic conditions [29–31] such as temperature and precipitation [27,32,33]. Changes in phenology can have community-wide consequences, and differential responses among various species within a community can lead to trophic mismatches [28,30]. For example, the timing of larval winter moth (*Operophtera brumata*) emergence was formerly largely synchronized with oak (*Quercus robur*) bud burst. Caterpillars that emerge too early lack a sufficient food source and will starve, while caterpillars that emerge too late will be exposed to older, poor-quality leaves, leading to negative physiological implications [34], Increased spring temperature has resulted in changes in the timing of oak bud burst. However, the winter moth has yet to adapt to changing temperatures, which has led to disrupted synchrony between these two species [34], Thus, phenological shifts can have both top-down and bottom-up consequences extending throughout multiple trophic levels. Long-term observations are important for understanding ecological trends and the merit of phenology as a predictor of ecological consequences. A long-term study on the Genji firefly (*Luciola cruciate*) in Japan found that populations patterns changed in response to rainfall, potentially leading to early larval emergence and reduced foraging [35]. However, the ways in which climate change and other environmental events have impacted firefly species is less understood.

Developing a model for the emergence of adult fireflies is key to developing our understanding of firefly phenology, which can then be used to expand firefly conservation efforts, educational outreach, environmental research, and to predict peak firefly display. In this study, we examine a ‘bycatch’ dataset documenting captures of fireflies at the Kellogg Biological Station over a 12-year period and place it in the context of other available data to gain insights into the long-term dynamics and phenology of this charismatic, but understudied taxon.

## Methods

### Data sources

Data were obtained through two publicly available data sets—a weather data set that included daily maximum and minimum temperature and precipitation, as well as a data set that focuses on ladybeetle observations, but also documents captures of the other insect species. Both data sets arise from Michigan State University’s Kellogg Biological Stations (KBS), located in southwestern Michigan. The firefly abundance data were collected as a part of the KBS Long Term Ecological Research (LTER) site within the Main Cropping System Experiment (MCSE) and forest sites starting in 2004. Fireflies were recorded to family alone, however, from spot-checks of the collected data, it appears fireflies collected belonged to the genus *Phot’mus*, mainly *P. pyralis*, the big dipper firefly, although captures of other species cannot be excluded.

Within the MCSE, seven plant community treatments were established in 1989, ranging from a three-year rotation of annual field crops (maize, soybean, wheat) under four levels of management intensity (conventional, no-till, reduced input, or biologically based), to perennial crops including alfalfa, poplar and early successional vegetation (*i.e.* abandoned agricultural fields maintained in an early successional state by yearly burnings; Table 1, Figure 1). Each of these treatments is replicated six times across the MCSE site with each replicate consisting of a 1 ha plot. We also included three forest sites in our analysis; these sites were established in 1993 within 3 km of the MCSE site on KBS and represent one of three plant community treatments: conifer forest plantations; late-successional deciduous forest; and successional forest arising on abandoned agricultural land (Table 1, Figure 1). Forested treatment plots are also 1 ha in size but are replicated three times for each treatment.

**Figure 1:**
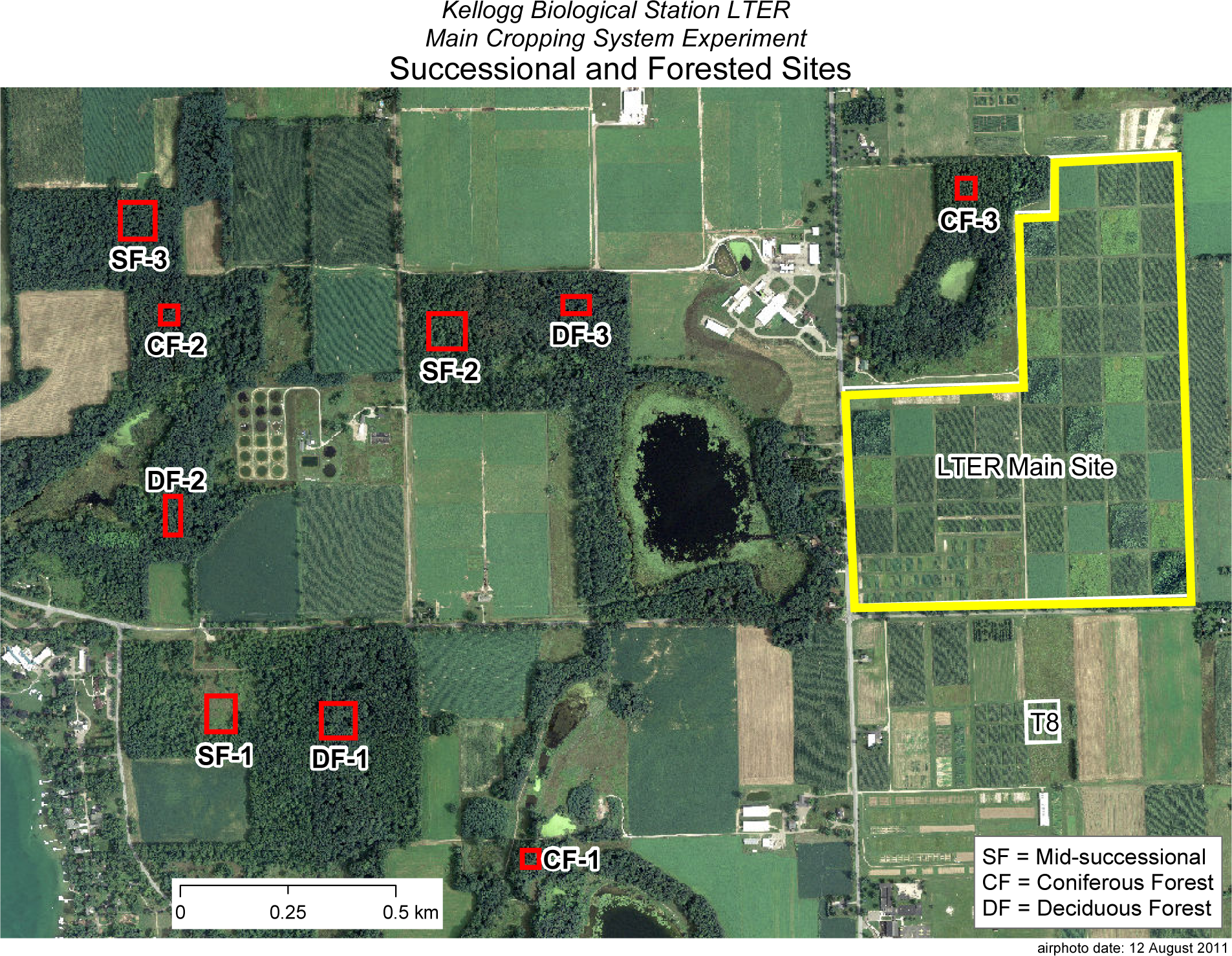
Map of Sites at the Kellogg Biological Station LTER. Site map of the Kellogg Biological Station Long-term Ecological Research Site (reproduced and modified from http://lter.kbs.msu.edu/maps/images/current-lter-forest-successional-sites.pdf). Red outlined areas indicate successional and forested sites, denoted by the key on the bottom right of the photo. Outlined in yellow is the main cropping system experiment (LTER Main Site) which houses 7 treatments: conventional crop, no-till crop, reduced input crop, biologically based crop, poplar, alfalfa, and early successional. Each plot is 1 ha and were replicated 6 times each.

**Table 1.**

| Crop Type | Plant Community Treatments | Description |
| --- | --- | --- |
| Annual | Conventional | Rotated crop field (corn-soybean-wheat), with conventional chemical input which is chisel plowed |
|  | No-Till | Rotated crop field (corn-soybean-wheat), with conventional chemical input, with no tilling. |
|  | Reduced Input | Biologically based rotated crop field (corn-soybean-wheat), with low input chemical control and a winter cover crop (leguminous). Plots are treated with banded herbicide and starter N at planting |
|  | Organic | Biologically based rotated crop field (corn-soybean-wheat), low input chemical control with winter cover crop (leguminous). Certified Organic |
| Perennial | Poplar Trees | 10-year rotation cycle of a fast growing *Populus* clone |
|  | Alfalfa | Continuously grown alfalfa |
|  | Early Successional | Abandoned field from 1989, left to grow into native successional plants which are annually burned |
| Forest | Successional | 40-60 year old successional forest, left from former agricultural fields |
|  | Coniferous | 40-60 year old conifer plantations |
|  | Deciduous | Late successional deciduous forest |

Observations were taken on a weekly basis throughout the sampling season at five sampling stations within each replicate (both MCSE and forest sites). These insect abundance data are available publicly, online at http://lter.kbs.msu.edu/datatables/67. Insect abundance monitoring was done using un-baited two-sided yellow cardboard sticky cards (Pherocon, Zoecon, Palo Alto, CA) suspended from a metal post within each sampling station, 1.2m above the ground. Cards were deployed each week for a one-week exposure for the duration of the growing season. Sampling start and end dates varied each year depending upon planting date of the various crops, however, the length of sampling was fairly consistent (14 ± 1 weeks, on average, per year).

In addition to plant community treatment management information, we also included weather as an environmental factor to explain firefly abundance. These data were also obtained through a publicly available data set, online at http://lter.kbs.msu.edu/datatables/7.

### Data pre-processing

All analyses performed are available as an R script at https://github.com/cbahlai/lampyrid/. Analyses presented in this manuscript were run in R 3.3.1 “Bug in Your Hair” (R Development Core Team 2016). Firefly data were extracted from the database held at the KBS data archive and combined with relevant agronomic data (which are encoded in plot and treatment numbers in the main database) and are hosted at figshare at https://ndownloader.figshare.com/files/3686040.

Data were subject to quality control manipulations to remove misspellings in variable names that had occurred with data entry. Observations with missing values for firefly counts were excluded from analysis. Because subsample data were zero-biased, we used reshape2 [36] to sum within date, within plot numbers of captures, and created an additional variable to account for sampling effort (which was usually consistent at five traps per plot per sampling period, but on occasion traps were lost or damaged).

Weather data (daily maximum and minimum temperatures were reported in °C and daily precipitation in mm) were downloaded directly from the Kellogg Biological Station Data archive (http://lter.kbs.msu.edu/datatables/7.csv). To overcome errors in calculations requiring accumulated annual weather data caused by rare missing data points (most often occurring during winter, in periods of extreme cold leading to equipment malfunction), we created a function to replace missing values in the temperature data with the value that was observed for that variable from the day before the missing observation.

We created a dummy variable for ‘start day’ to enable the user to test the sensitivity of our conclusions to varying our within-year start of accumulation of environmental conditions. We empirically determined (By AIC) that March 1 (start = 60) provided the best compromise between capturing early growing season weather variation and negating brief variation in winter conditions, however, the selection of the precise day did not dramatically influence the overall trends in the results unless changed by >15 days.

We then created a function to calculate daily degree day accumulation and season-long degree day accumulation based on Allen’s [37] double sine function, using our daily maximum and minimum temperature data. We created a dummy variable for our minimum development threshold to facilitate sensitivity analysis, but set it to a default value of 10°C. We did not use a maximum development threshold in our calculation, assuming that temperatures exceeding its hypothetical value (often >30°C for temperate insects) were relatively rare. Accumulations were calculated from the start day variable, as described above. We also created functions that calculated the accumulation of precipitation over the sampling week, the accumulation of precipitation over the growing season, from the start date, and the number of rainy days in a sampling period. Weather data were merged with firefly data to facilitate subsequent analyses.

### Data analysis

We used ggplot2 [38] to visualize trends in captures of fireflies by plant community treatment over years. We then conducted a multivariate analysis to determine if firefly plant community use patterns changed within or between years, and what environmental factors were associated with plant community use patterns. To accomplish this, data were cast as a date-by-treatment matrix at two resolutions (weekly observations and yearly observations), transformed using the Wisconsin standardization, and Bray-Curtis differences were subjected to non-metric multi-dimensional scaling (NMDS) in vegan [39]. Environmental parameters were fit to the NMDS plots using envfit to determine if patterns were influenced by weather.

To examine patterns in firefly captures over time, and the interactions of these captures with environmental variables, we visualized trends in capture data by sampling week and degree day accumulation. Noting that degree day accumulation was associated with the clearest patterns in firefly captures (see results), with some variation due to plant community, we built a generalized linear model with a negative binomial structure to explain these patterns. The model included the degree day accumulation in linear and quadratic forms as continuous variables, year and plant community treatment as factors, and trapping effort as an offset variable (to account for lost or compromised traps). Model structure was determined empirically by AIC. After fitting the model, we used the resultant regression parameters to generate predicted values, so we could visually compare the performance of the model to the raw data.

Because the model found year-to-year variation in the activity peak that was not explained by degree day accumulation, we extracted the activity peaks from each year as predicted by the model to a new data frame, and matched these data to other relevant environmental variables in the weather matrix (week the peak occurred in, precipitation variables corresponding to that week). We visualized the relationship between activity peak and other variables, and then constructed a generalized linear model for a quadratic relationship between the activity peak, by degree days, and the precipitation accumulation at the activity peak.

For all frequentist analyses, a significance level of α = 0.05 was used.

## Results

Over the 12 year study, 17 084 fireflies were captured in the trapping network. Visualizations of firefly capture data by treatment and time period revealed several patterns. Numbers of fireflies captured in each trap varied by plant community type and across samples (Figure 2), but in general, more fireflies were captured in alfalfa and no-till row crop treatments. Average numbers of fireflies captured per trap also demonstrated variation by year independent of plant community treatment. Overlaid plots of average captures for all treatments against year (Figure 3) suggest a 6-7 year firefly population cycle that appears uncorrelated with environmental variables.

**Figure 2:**
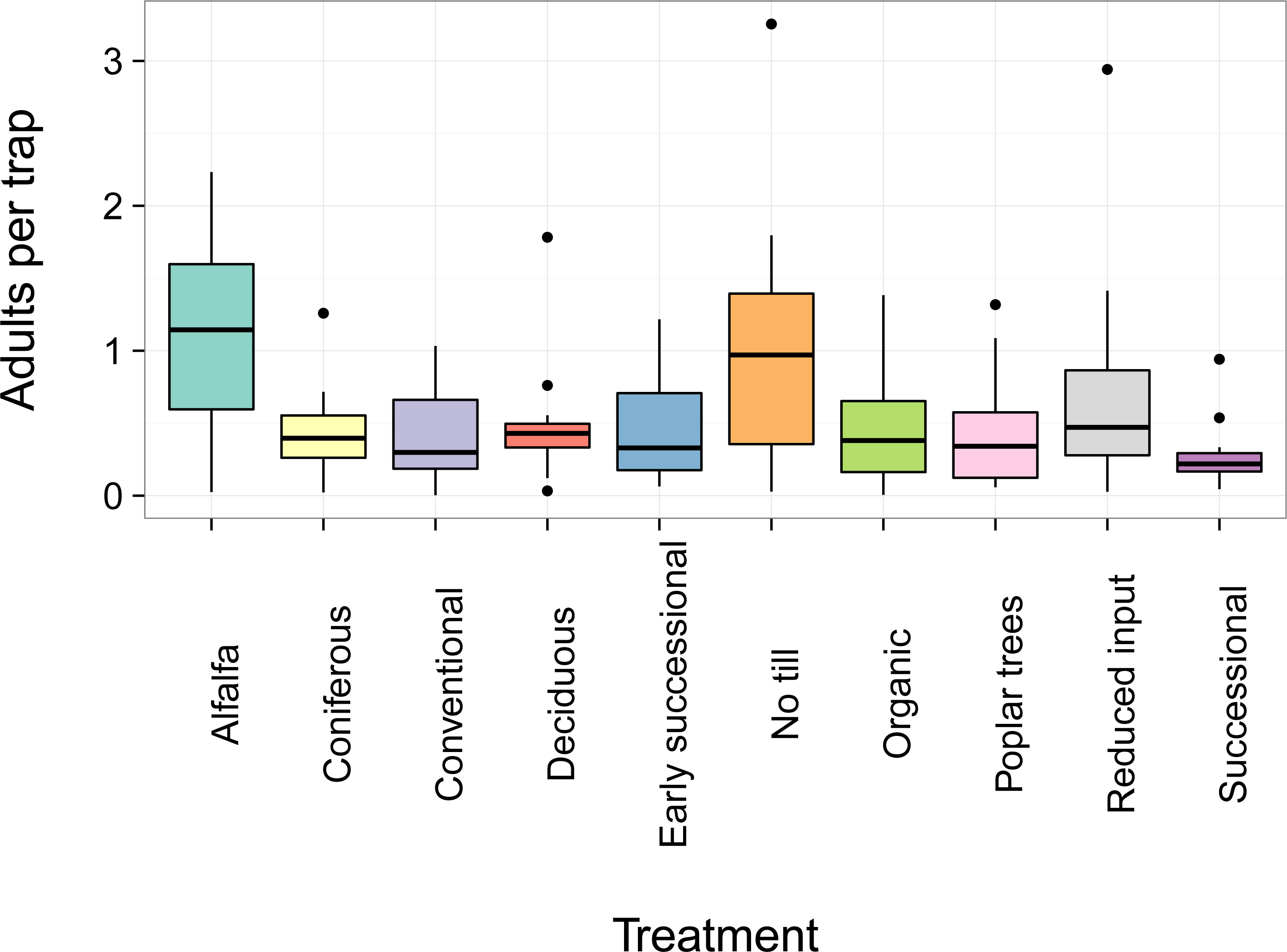
Box plot of average firefly captures, 2014-2015, by plant community treatment. Yearly average number of adult fireflies captured on weekly sampled yellow sticky cards across ten plant community treatments at Kellogg Biological Station. Median firefly density in each treatment is represented by the bold line, and upper and lower margins of each box represent the upper and lower quartiles in that treatment, respectively.

**Figure 3:**
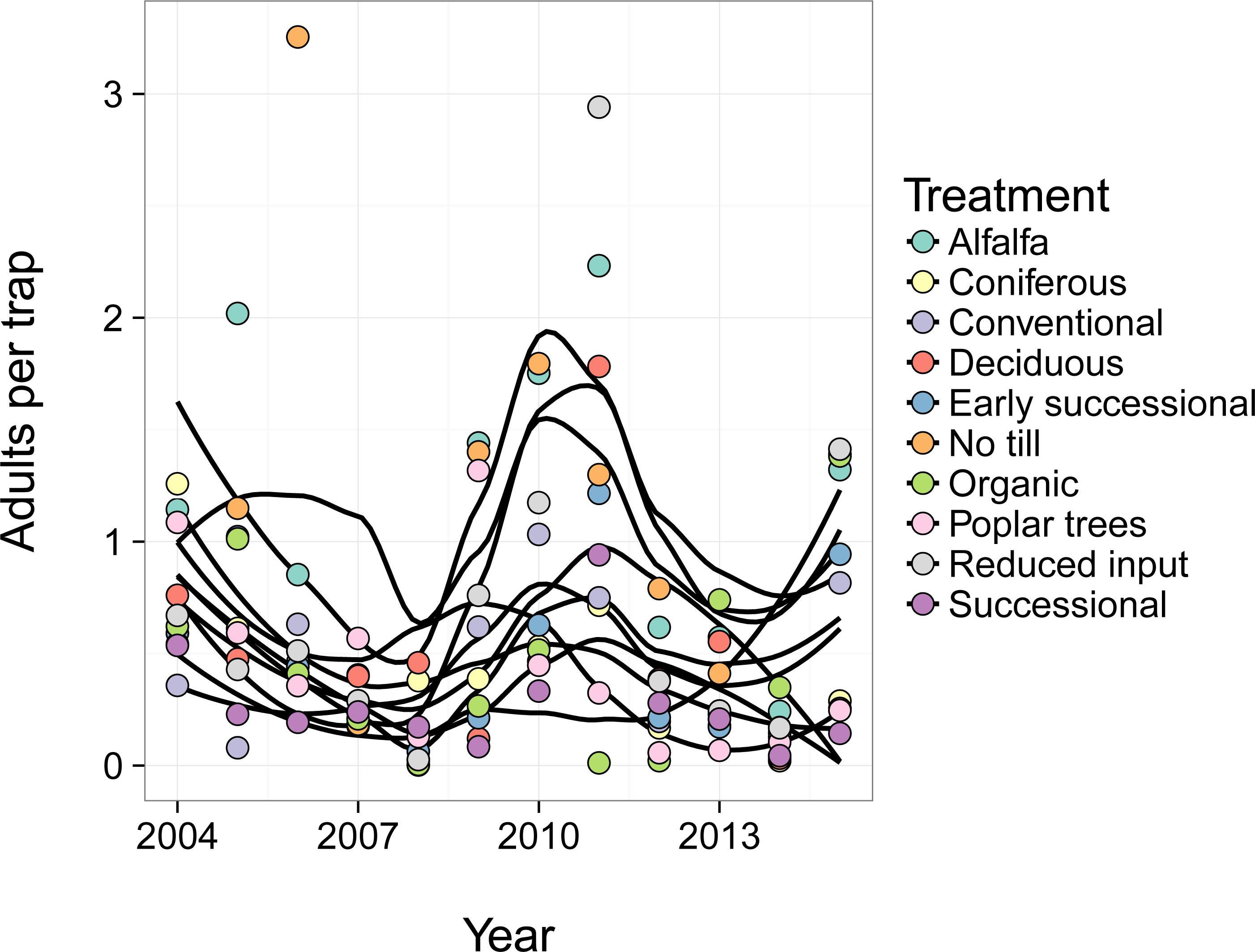
Average firefly captures, 2004-2015, by plant community treatment, by year. Yearly average number of adult fireflies captured on weekly sampled yellow sticky cards across ten plant community treatments at Kellogg Biological Station. Loess smoother lines represent smoothed captures within a given treatment and are used to illustrate general trends in population across treatments.

Non-metric multi-dimensional scaling revealed only weak trends in patterns of capture between plant community treatments at both the yearly and weekly resolutions. At the yearly resolution (Figure 4A), plant community treatment use varied slightly with the number of rainy days in the growing season (R^2^ = 0.16, p = 0.006, 2-D NMDS stress = 0.14) with herbaceous habitat use generally associated with greater amounts of rainfall. At the weekly resolution (Figure 4B), 2D-NMDS stress was higher (0.19), but a general trend away from forest plots was observed with increasing degree day accumulation (R^2^ = 0.15, p = 0.001) and week (R^2^ = 0.15, p = 0.001).

**Figure 4:**
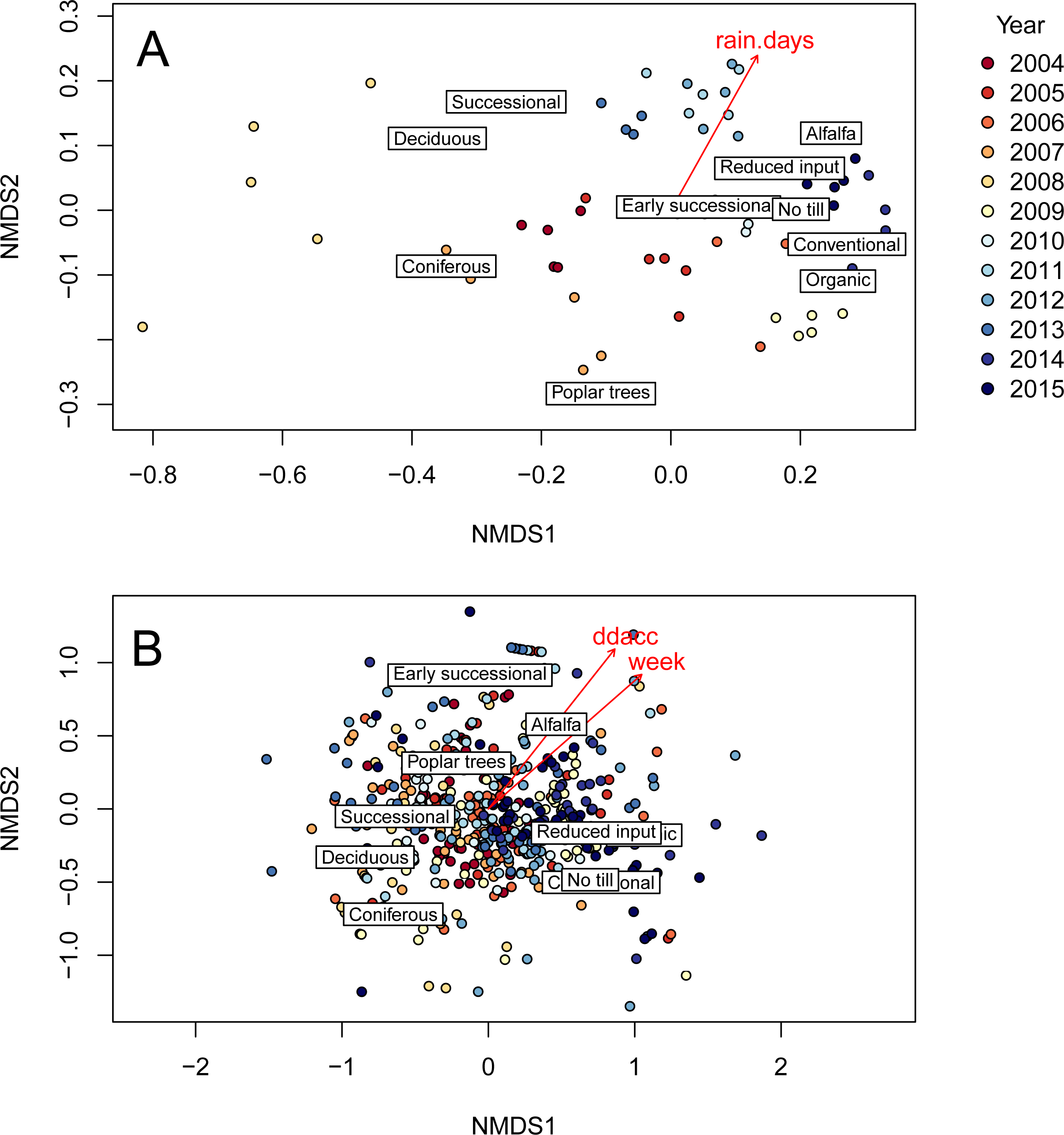
Two-dimensional non-metric multidimensional scaling and environmental fitting of plant community treatment plot use by fireflies over time. A) At the yearly resolution, a 2D NMDS stress of 0.14 was observed. B) At the weekly resolution, a 2D-NMDS stress of 0.19 was observed.

When plotting firefly abundance by week of capture, the timing of the peaks in firefly emergence show asynchrony among years (Figure 5A), indicating that week of year (and, by proxy, day length) is not a strong driver of firefly emergence. However, plotting firefly numbers instead against degree day accumulation dramatically reduced the asynchrony of emergence peaks and indicated a single activity peak occurred in each year (Figure 5B). Thus, degree day accumulation appears to be a better predictor of firefly populations than week of year or associated variables.

**Figure 5:**
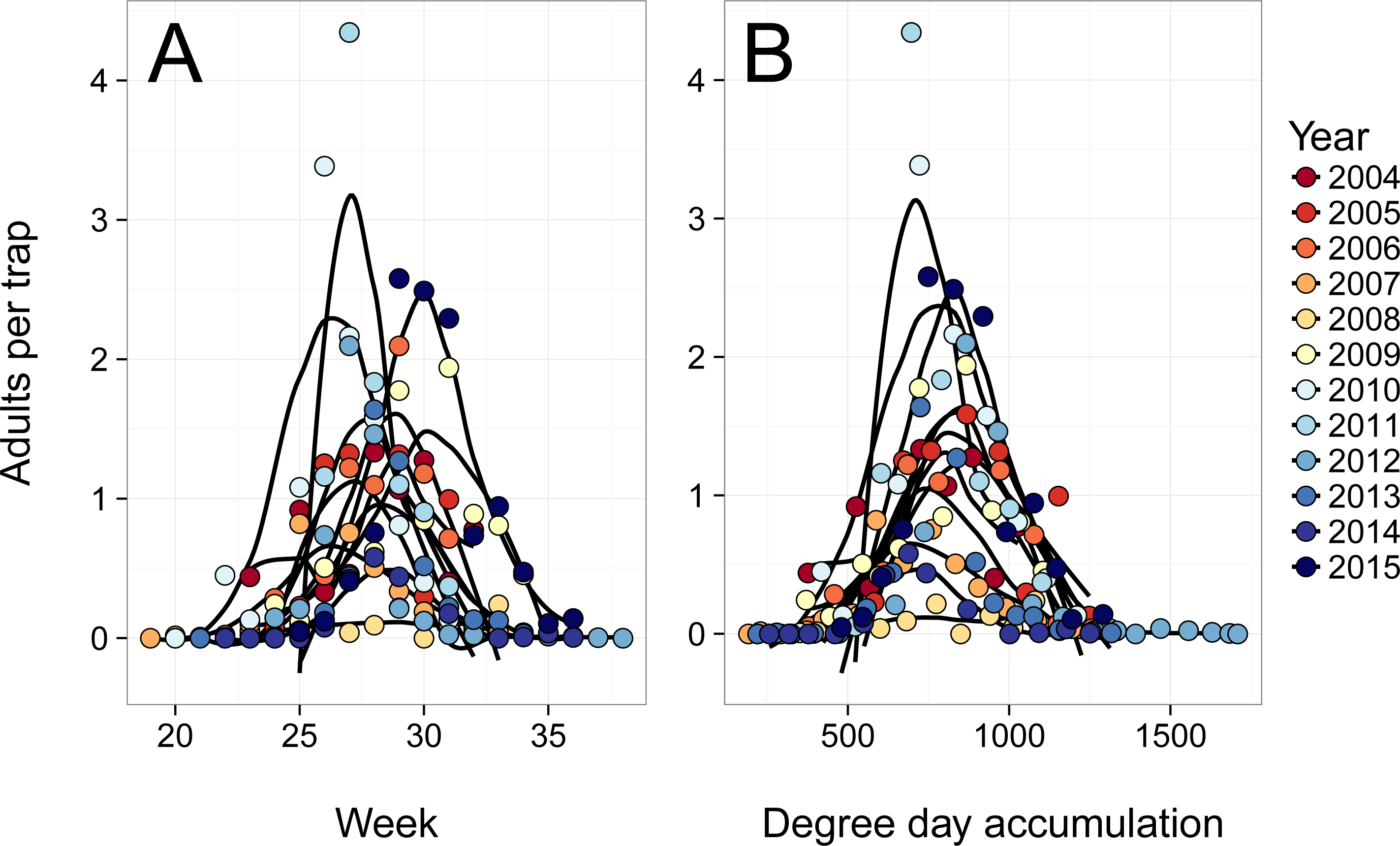
Average number of adult fireflies per trap across all sampled treatments at Kellogg Biological Station plotted by year. Samples were taken weekly over the growing season from 2004-2015, and plotted by A) week of capture; and B) degree day accumulation at capture. Loess lines represent smoothed capture trends for a given year and were used to assess consistency of response to a given variable between years.

Our model for firefly activity incorporating degree day accumulation, plant community treatment, and year, performed well at predicting the timing of the activity peaks (Figure 6), accounting for more than 40 percent of the variation in the raw data. However, model selection favored the inclusion of a year term as a factor, suggesting that another factor in addition to degree day accumulation was varying from year to year, and impacting firefly activity. Activity peaks varied from year to year by nearly 180 degree-day units, varying from 720 ± 38 DD in 2004 to 898 ± 55 DD in 2012 (Figure 7). However, we found the year-to-year variation was well-explained by precipitation accumulation: a quadratic relationship occurs between degree day at peak emergence and precipitation accumulation (pseudo-R^2^ = 0.456, *p* = 0.026; Figure 8).

**Figure 6:**
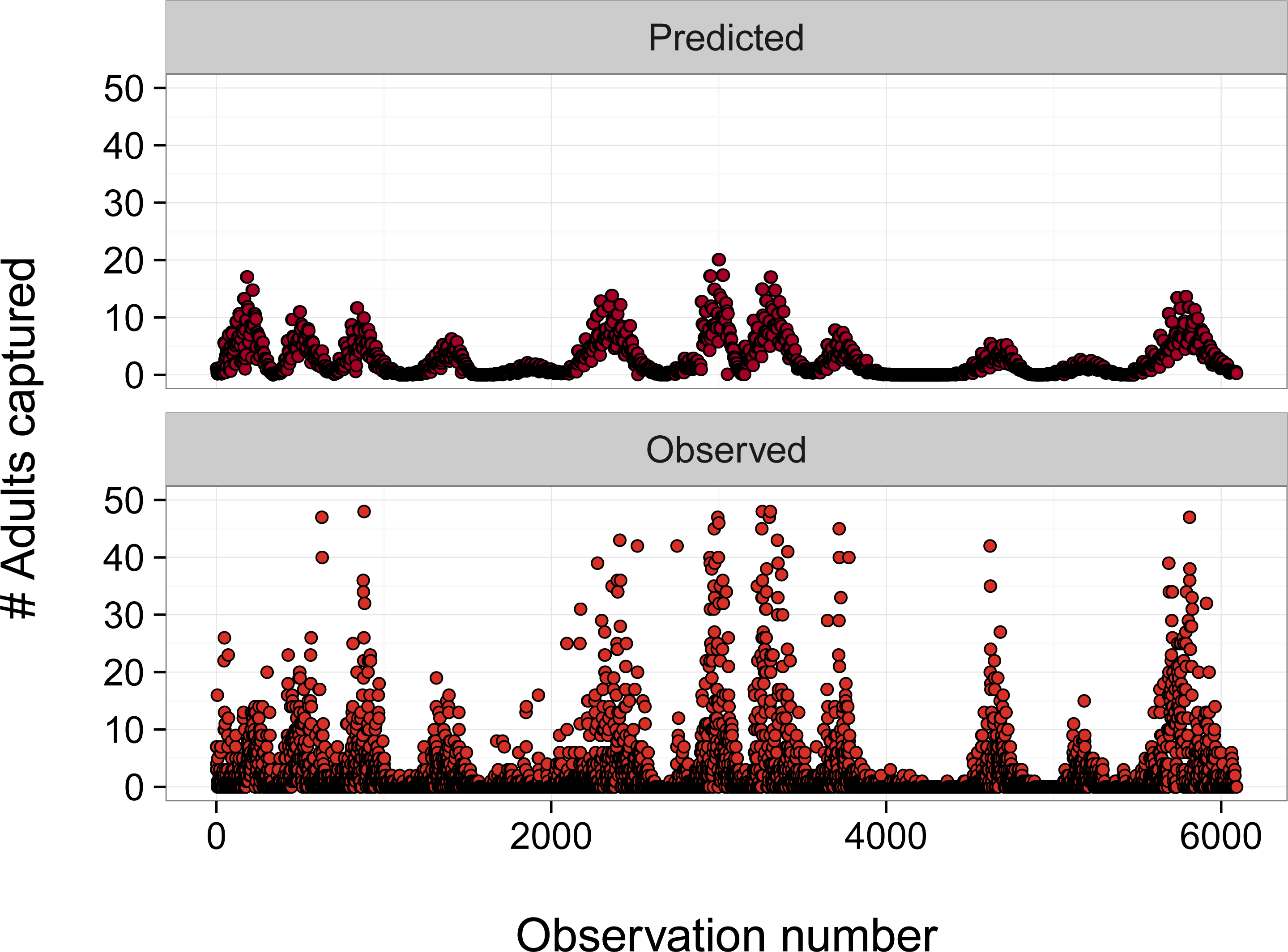
Number of firefly adults captured, as predicted by GLM and as observed, by observation number. Predicted values were generated using GLM accounting for variability due to plant community treatment degree day accumulation, and year as a factor variable. Details of GLM can be found in data analysis section in Materials and Methods.

**Figure 7:**
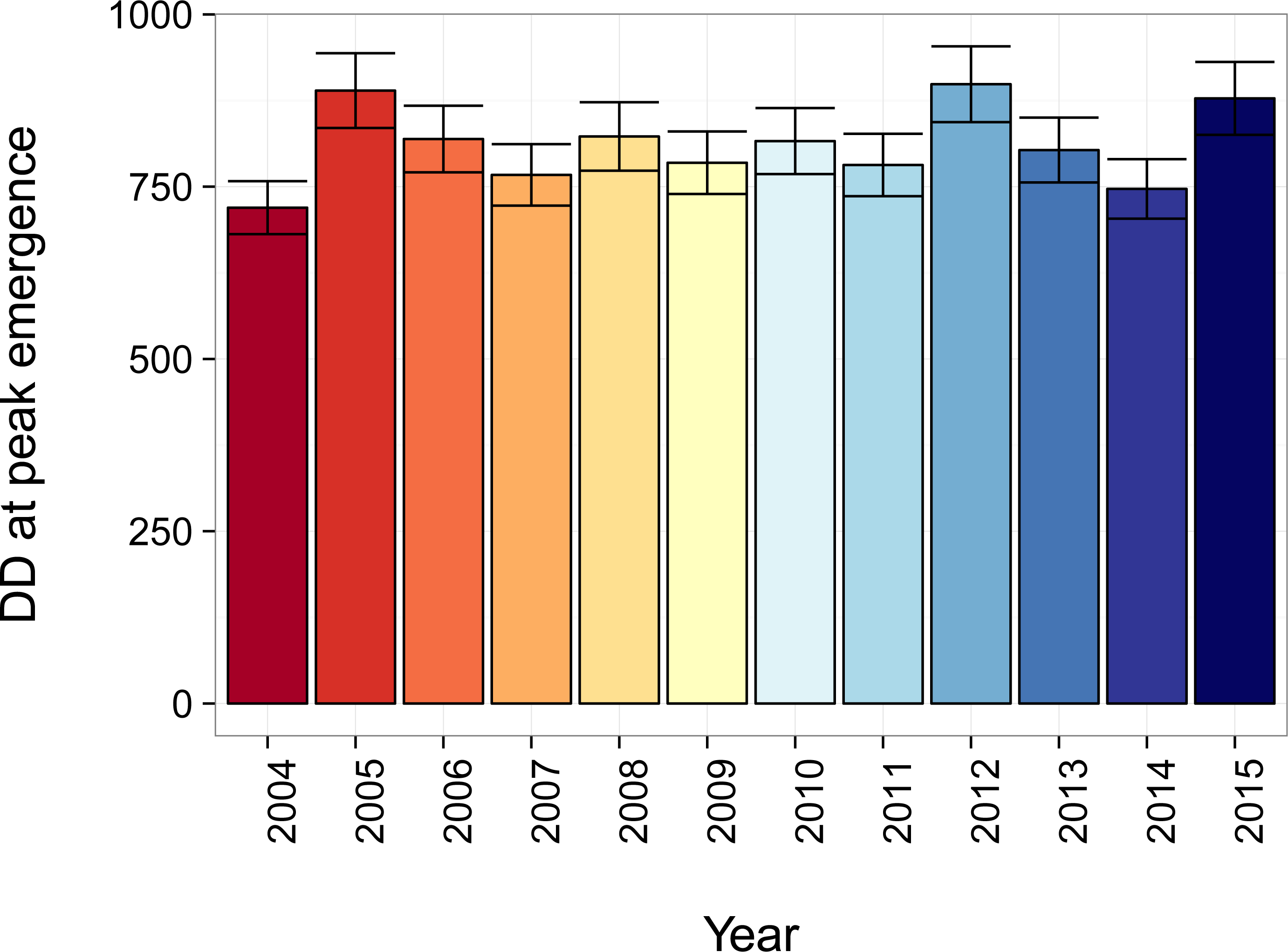
Degree day accumulation at peak firefly activity by year. Degree day accumulation (±SEM) at peak emergence of firefly adults varied by sample year. Activity peaks were extracted from regression coefficients from GLM.

**Figure 8:**
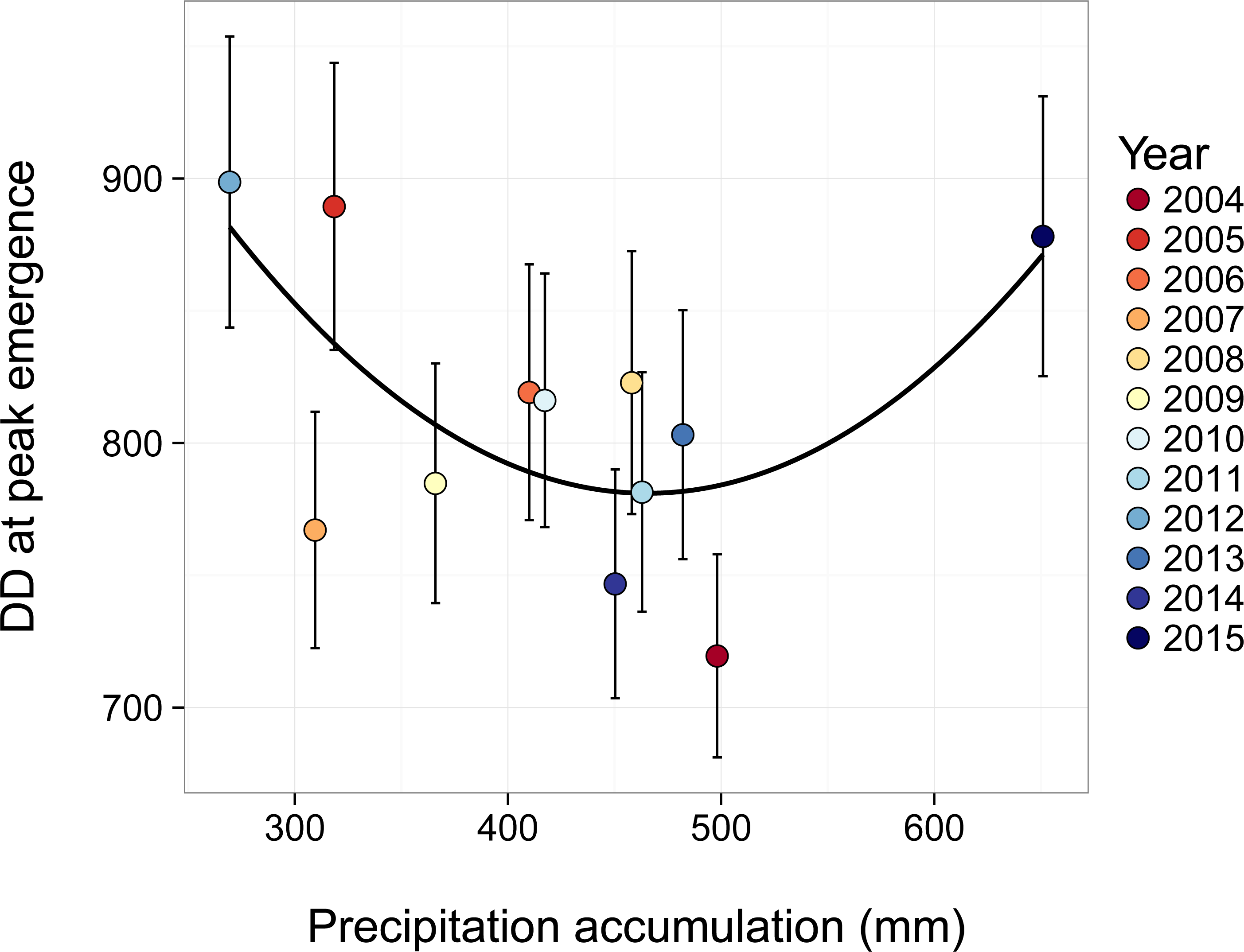
Firefly activity peaks by precipitation accumulation. Firefly activity per degree day accumulation had a quadratic relationship with precipitation accumulation (pseudo-R^2^= 0.456, p = 0.026).

## Discussion

The greatest proportion of fireflies was captured in alfalfa and no-till plant communities (Figure 2), indicating that areas with moderate soil disturbance and primarily herbaceous plant communities favored firefly emergence. This result was unexpected; because fireflies spend much of their life cycle in the soil, it might be expected that plots with little soil disturbance (coniferous, deciduous and successional forests) would foster the greatest populations of fireflies. However, these plots produced capture rates similar to those observed in the intensively managed and tilled conventional row crop plots. Our result contrasted with observations of another genus of fireflies in Malaysia (*Pteroptyx*), where researchers found that plant canopy structure was the most important determinant of abundance [40]. Also surprising was the relatively low capture rate in early successional plots, which are primarily herbaceous, with no tillage regime. Thus, the yearly burnings may play a role in suppressing firefly populations in these plots. An alternative explanation for these variations in captures could be differences in trapping efficiencies between plant communities. However, if this were the case, we would expect trapping efficiencies in the three other row crop treatments (conventional, organic and reduced input management) not to differ appreciably from that of the no-till row crop plant community.

When plotted over sample years (Figure 3), captures of fireflies by treatment seem to suggest an intriguing cyclical dynamic, with alternating peaks and troughs in captures on an approximately 6-year cycle. Our time series only spans 12 years, meaning more data will be required to elucidate this pattern and its drivers. Similarly, analysis of plant community use patterns was inconclusive (Figure 4). At the weekly resolution there was a trend away from woody treatments over the growing season (Figure 4B; *i.e.* with both increasing week and increasing degree day accumulation). Although this pattern was not strong, it could result from fireflies overwintering in forest habitats and then moving to lower-canopy herbaceous habitats for mating displays. We observed very similar performance of both degree day and week, likely due to autocorrelation between the two variables that cannot be resolved at the sampling resolution used over the course of the study.

The degree-day model GLM suggested activity peaks occurred at a degree day accumulation of ~800 DD, accumulated from March 1. A model for *Photinus carolinus* in the Great Smokey mountains found peak display occurred at ~1100 degree days (using a base of 10°C and the same start date as our model) [25]. The difference between heat units required for peak activity observed between this and our study may be a result of species or locality differences (i.e. more southern firefly populations are likely adapted to warmer spring conditions). Similarly, differences in methodology for calculating degree days may account for some of these differences. However, both studies support the observation that degree day accumulation is the dominant cue governing the activity patterns of temperate fireflies.

Although both photoperiod and degree day accumulation can both play a role in the phenology of insects, our results suggest that degree day accumulation is the dominant driver of firefly flight activity. The model was unable to account for between-trap variation within a single sampling day (Figure 6), though it was able to capture the overall trends in activity quite well, using only degree day accumulation, plant community treatment, and year as predictors. Nevertheless, degree day accumulation was not the sole driver in within-season variability. Our model found year-to-year variability in activity peaks that could not be explained by degree day accumulation alone. We found that this variation in activity peak by degree day accumulation had a quadratic relationship with precipitation, indicating that both drought and heavy rainfall in the time period leading up to their activity peak can delay the peak (Figure 8). Assuming a ~20°C daily average temperature at this site in late June and early July, this could translate to a ten-or-more day change in activity peak due to precipitation extremes in any given year. Yet, there are several alternate explanations for the pattern we observed, and some patterns detected may have been driven by statistical outliers. For example, very high rainfall in 2015 at the site strongly influences our conclusion that a quadratic relationship exists between degree day accumulation and precipitation accumulation in explaining firefly activity peaks. Indeed, if observations from 2015 had not been included in our analysis, we would likely have concluded that degree day accumulation had a negative, linear relationship with precipitation accumulation, that is, increasing rainfall would cause fireflies to emerge earlier, given constant temperatures. This result would align with previous work showing firefly abundance in Japan is generally negatively correlated with rainfall [35]. However, considered within the context of firefly biology, it seems unlikely that the relationship between these parameters would be linear throughout the rage of possible precipitation values, as soil dwelling larvae of non-aquatic firefly species and/or their prey are likely adversely affected by abnormally waterlogged soils.

As the sampling at our study site continues, we will watch rainy years with particular interest to determine if population data collected in these years support or refute this pattern, or if an alternate driver can explain more of the variation. Indeed, firefly activity may have been driven by factors not considered in this study. Although using a start date of March 1 was favored in our analysis (i.e. the AIC of the model using this start date was minimized), when the start day was changed in a sensitivity analysis, the relationship between degree day accumulation and precipitation in firefly activity changed or disappeared. This result could suggest that alternate drivers not accounted for in this study may be driving aspects of firefly activity. Factors such as winter snow cover and variations in winter temperature are known to affect the phenology of temperate insects [41], and thus these factors should be considered in subsequent work.

In this study, we have clearly demonstrated a taxon whose phenology varies in response to multiple drivers. Species with phenological responses to multiple drivers are not rare [42]. Yet ecological interactions among species with multiple drivers of phenology may be complex and unpredictable [43,44], potentially leading to dire consequences in a changing environment [45]. Our study examined the phenological responses to environmental conditions of adult fireflies; however, data on larvae or sex of the adults were unavailable. Adult *Photinus* fireflies are non-feeding [14], so shifts in their activity are unlikely to have direct consequences through phenological asynchronies. Shifts in adult activity likely correspond to shifts in development or activity among larvae, potentially leading to asynchronies between larvae-prey populations at this critical development time-period. Resources acquired during the predaceous larval stage are important in determining mating success among adult fireflies: males provide an energetically costly nuptial gift to the female in the form of a spermatophore [46]. If sex differences in phenological responses to environmental conditions exist, asynchronies between males and females may additionally reduce mating success and fecundity [47]. Male fireflies were always observed earlier than females in Elkmont, Tennessee, USA. In fact, in that system, females were often found during or after peak emergence of males and thus this should be an area of emphasis in future study [25]. Additionally, phenological shifts in fireflies may lead to consequences at other trophic levels. For example, generalist ground-dwelling predators like firefly larvae and other predaceous beetles are known to have dramatic effects on the establishment of agricultural pests early in the growing season [48]. Similarly, although distasteful and avoided by many predators, some birds, lizards and frogs are known to feed on adult fireflies [49], thus shifts in firefly activity may have dietary consequences for animals at higher trophic levels.

## Conclusions

Fireflies are a charismatic and important taxon with ties to trophic function, economic importance, and culture. Although empirical evidence of specific declines of *Photinus* fireflies has not been clearly demonstrated in longitudinal studies, naturalists and citizen scientists perceive a decline in their number [21], leading to interest in their conservation. Our study has offered new insight to support conservation efforts and to direct future research. *Photinus pyralis* appears to thrive in habitats with moderate soil disturbance. Thus, efforts to foster no-till and perennial agricultural systems [50,51] will likely benefit the species. Climate warming may advance the activity of fireflies to earlier and earlier in the growing season, but other extremes of climate in the form of precipitation may introduce unpredictable elements to this, and add the possibility of inducing asynchrony with other systems.

The availability of long-term observational data, made freely accessible to the public, was an essential factor in the discoveries made in this study. Although the study that provided these data was not initiated with this purpose in mind, we were able to empirically demonstrate and disentangle the effect of multiple drivers on firefly phenology simply because we had the statistical power to do so. Although species that respond to multiple, interacting environmental drivers are relatively common, data supporting investigations of this kind are rare [52]. We therefore encourage all practicing ecologists to curate their species observation data and make them publicly available, to foster long-term, broad-scale investigations in the future [53–55].

## Research Ethics

This study did not require the approval of an ethics committee.

## Animal Ethics

This study did not use animal subjects.

## Permission to carry out fieldwork

No new fieldwork was carried out by this study. Data used by this study were part of an ongoing NSF-funded field trial conducted on a field station owned by Michigan State University.

## Data Availability

All data and analysis code produced by this study are publicly available. Lampyrid abundance data are available at https://ndownloader.figshare.com/files/3686040. Weather station data are available at http://lter.kbs.msu.edu/datatables/7.csv. These data are automatically downloaded when the R script file at https://github.com/cbahlai/lampyrid/ is run.

## Author’s contributions

SH contributed to conception and led study design and wrote major portions of the manuscript. SX conceived of analyses and wrote portions of the manuscript. LR led exploratory analysis and wrote portions of the manuscript. EDL, AM and BE provided critical commentary in the design and analysis of the study and wrote portions of the manuscript. CB conceived of the study, led the analysis, supervised the drafting of the manuscript, and critically revised its content. All authors approved the final content of the manuscript.

## Acknowledgments

This paper was written as part of a course in Open Science and Reproducible Research offered through the Department of Entomology at Michigan State University. The authors would like to thank the Mozilla Science Lab and the rest of the open science community for their support in the drafting of this manuscript, and Stuart Gage, Manuel Colunga-Garcia and Douglas Landis for design and maintenance of the study that produced the data used herein. Monica Granados helpfully provided review of our analysis R code. We would like to thank Tyson Wepprich and one additional anonymous reviewer for helpful comments that helped us question our assumptions and improve the clarity of this manuscript.

